# GL-0001, a first-in-class dual GIP/GLP-2 analogue, reduces bone fragility by enhancing bone material properties in a murine model of type III osteogenesis imperfecta

**DOI:** 10.64898/2026.10.16.759580

**Authors:** Malory Couchot, Antoine Chretien, Aleksandra Mieczkowska, Catherine Behets, Guillaume Mabilleau

## Abstract

Osteogenesis imperfecta (OI) is a severe heritable skeletal disorder characterized by bone fragility, short stature, and increased fracture risk. Currently, no pharmacological treatment has been approved for OI, and bisphosphonates are commonly used off label. While bisphosphonates increase bone mass, their effects on bone quality and fracture reduction remain limited. GL-0001, a first-in-class dual glucose-dependent insulinotropic polypeptide (GIP)/glucagon-like peptide-2 (GLP-2) compound that targets bone material properties, may offer alternative skeletal benefits. We evaluated the effects of 9 weeks of GL-0001 treatment in the oim/oim murine model of severe OI, a well-established model carrying a homozygous *Col1a2* mutation resulting in abnormal type I collagen formation and compared outcomes with zoledronic acid (100 μg/kg) or saline. Analyses included longitudinal growth, fracture incidence, bone biomechanics, microarchitecture, turnover markers, and bone material properties. OI mice exhibited reduced body weight, impaired femoral growth, severe trabecular and cortical deficits, altered bone turnover, and compromised biomechanical properties compared with WT controls. GL-0001 increased femur length by 5% (p=0.015) and reduced spontaneous fracture incidence by 90% (p=0.004), whereas zoledronic acid had no significant effect on fracture burden. Bone microarchitecture was not significantly altered by GL-0001, whereas zoledronic acid improved trabecular and cortical parameters. At the tissue level, GL-0001 improved bone strength, increasing ultimate stress (+26%, p=0.022) and yield stress (+30%, p=0.014), and reducing brittleness (−32%, p=0.002). These changes were associated with increased collagen maturity (+12%, p<0.001). In contrast, zoledronic acid had limited effects on bone material properties. These findings demonstrate that GL-0001 improves skeletal fragility in the oim/oim murine model of severe OI through mechanisms primarily involving enhancement of bone material properties and longitudinal growth, rather than changes in bone mass or microarchitecture. GL-0001 may therefore represent a novel therapeutic strategy targeting bone quality in OI.

## 1. Introduction

Osteogenesis imperfecta (OI) is a heritable connective tissue disorder characterized by skeletal fragility, bone deformity, and impaired longitudinal growth, with an incidence of approximately 1 in 10,000 to 15,000 births. Approximately 85% of patients harbor pathogenic variants in *COL1A1* (OMIM 120150) or *COL1A2* (OMIM 120150), which encode the pro-α1(I) and pro-α2(I) chains of type I collagen, respectively ^(1)^. These chains assemble in a 2:1 stoichiometry into a triple-helical heterotrimer that forms collagen type I fibrils ^(2)^. Type I collagen is the principal organic component of bone, providing tensile strength and serving as a scaffold for hydroxyapatite deposition, thereby contributing to bone mineralization and mechanical competence ^(3)^. OI is clinically heterogeneous, ranging from mild osteopenia to severe skeletal deformities. Type III OI, according to the Sillence classification, is characterized by severe short stature, progressive skeletal deformities, and early-onset recurrent fractures. These phenotypic features reflect profound alterations in both bone quantity and bone quality, including abnormal collagen matrix organization, reduced bone mass, impaired microarchitecture, and compromised tissue-level mechanical properties ^(4)^. The oim/oim mouse represents a well-established model of moderate-to-severe type III OI. This model carries a homozygous mutation in *Col1a2* resulting in a truncated α2(I) chain and formation of abnormal collagen type I homotrimers instead of the normal α1(I)_2_α2(I) heterotrimer ^(5)^. The oim/oim phenotype recapitulates key features of human OI, including spontaneous fractures and marked skeletal fragility associated with defects in bone mass, microarchitecture and material properties ^(6,7)^.

There is currently no approved curative therapy for OI, although intravenous and oral bisphosphonates are widely used in both pediatric and adult clinical practice. These agents consistently increase bone mineral density (BMD) compared with placebo; however, evidence for a corresponding reduction in fracture risk remains limited and inconsistent across studies ^(8,9)^. This apparent dissociation between gains in bone mass and fracture risk highlights fundamental differences between OI and osteoporosis, despite both conditions being characterized by increased skeletal fragility. Although the repositioning of osteoporosis drugs for osteogenesis imperfecta is an attractive idea, current evidence suggests that increasing bone mass alone may be insufficient to substantially reduce fracture risk in OI. Teriparatide has been investigated in adults with OI and was shown to increase BMD compared with placebo, although the study was not powered to assess fracture outcomes ^(10)^. Likewise, sclerostin inhibition has emerged as a promising anabolic strategy. In the phase 2b ASTEROID study, setrusumab produced dose-dependent increases in BMD over 12 months; however, no significant differences in annualized fracture rates were observed between treatment groups ^(11)^. Similarly, the ORBIT, COSMIC, and TOPaZ trials evaluating setrusumab or sequential teriparatide followed by zoledronic acid have demonstrated improvements in BMD without clear reductions in fracture incidence ^(12,13)^. Collectively, these clinical data suggest that improving bone mass alone may be insufficient to substantially reduce fracture risk in OI, particularly in severe forms of the disease where qualitative defects in type I collagen compromise bone material properties. This limitation has shifted attention toward therapeutic strategies that target bone quality, including matrix organization, collagen maturation, and tissue-level mechanical competence rather than solely bone density.

In this context, incretin-based pathways have recently emerged as regulators of skeletal tissue quality. Beyond their metabolic effects, glucose-dependent insulinotropic polypeptide (GIP) and glucagon-like peptide-2 (GLP-2) signaling have been shown to improve bone material properties and enhance collagen maturity in vitro and in vivo ^(14-17)^. These observations are particularly relevant in OI, where abnormalities in enzymatic collagen crosslinking and extracellular matrix organization have been documented across several genotypes ^(18-22)^. Accordingly, a new class of unimolecular dual GIP/GLP-2 analogues has been developed with the objective of restoring enzymatic collagen crosslinking and improving extracellular matrix quality. We therefore hypothesized that this approach may be particularly relevant for OI, since altered collagen organization is a primary determinant of bone fragility. Among these compounds, GL-0001 emerged as a particularly attractive candidate for OI because, in a model of estrogen deficiency–induced bone fragility, it improved bone biomechanics more effectively than zoledronic acid and restored compromised collagen maturity through a cAMP-lysyl oxidase-dependent mechanism ^(23)^.

Here, we investigated the effects of GL-0001 in the oim/oim murine model of type III OI and compared its effects with those of zoledronic acid across multiple levels of skeletal organization, including bone growth, microarchitecture, biomechanics, bone material properties, and fracture incidence. We hypothesized that, by improving collagen maturation and bone material properties, GL-0001 would enhance skeletal strength and reduce fragility in severe OI.

## 2. Materials and methods

### 2.1. Animals

All procedures were conducted in accordance with the European Union Directive 2010/63/EU for animal experiments and the Belgian federal regulations for animal experimentation and were approved by the Institutional Animal Care and Use Committee of the Université catholique de Louvain (protocol #2020/UCL/MD/02). Heterozygous mice (strain B6C3Fe a/a-Col1a2^oim^/J; JAX stock #001815) were obtained from The Jackson Laboratory (Charles River Laboratories, L’Arbresle, France). Heterozygous breeders were intercrossed to generate homozygous oim/oim and wild-type (WT) littermates. Five-week-old homozygous oim/oim mice were included in the study. Genotyping was performed by PCR amplification of genomic DNA extracted from tail biopsies using the following primers: forward (5′–3′) GGCTTTCCTAGACCCCGATGCTTAG; reverse WT (5′–3′) GTCTTGCCCCATTCATTTGTC; and reverse oim (5′–3′) GTCTTGCCCCATTCATTTGTT. Fifteen mice were initially assigned to each treatment group. At 5 weeks of age, following baseline imaging, only mice exhibiting at least one fracture, fracture callus, or skeletal deformity were included in the study. Eligible mice were then randomized to one of three treatment groups for 9 weeks: saline (n = 13), zoledronic acid (100 µg/kg, n = 6), or GL-0001 (25 nmol/kg/day, n = 9). Saline and GL-0001 were administered once daily by subcutaneous injection, whereas zoledronic acid was administered as a single intravenous injection at the beginning of the treatment period. To ensure comparable handling across treatment groups, mice receiving zoledronic acid were handled daily throughout the study, in the same manner as mice receiving saline or GL-0001. Supplementary figure 1 represents a schematic of animal study. The number of fractures per mouse at inclusion did not differ significantly between treatment groups. WT littermates treated with saline (n = 10) served as controls. Only female mice were used, based on previous evidence of sex-dependent differences in the oim model, with females exhibiting more severe alterations in bone microarchitecture, biomechanical properties, and bone material properties ^(24,25)^. Mice were housed in social groups at 24°C under a 12 h light/12 h dark cycle and had *ad libitum* access to standard rodent chow (ref. A03; Safe Diets, Augy, France) and water. No adverse events were observed in any group. Body weight was measured weekly. Fluorochrome labeling was performed by subcutaneous injection of calcein (30 mg/kg) at 10 and 2 days prior to euthanasia. At necropsy, blood collection by intracardiac aspiration (~250 µl) was performed in EDTA-treated tubes. Blood samples were spun at 13,000 g for 15 minutes, aliquoted and stored at −80°C until use.

### 2.2. In vivo imaging

#### 2.2.1. Acquisition/Reconstruction of in vivo CT images

In vivo imaging was performed using a NanoSPECT/CT PLUS system (Mediso, Budapest, Hungary) under isoflurane anesthesia. Imaging sessions were acquired at baseline (5 weeks of age, prior to treatment initiation), 4 weeks after treatment initiation, and at study endpoint (14 weeks of age). CT acquisitions were performed in helical mode using an X-ray tube voltage of 65 kV, a tube current of 120 µA, and an exposure time of 1000 ms per slice. An isotropic voxel size of 0.221 mm was used. The total scan time was <15 min per animal. The scanner was calibrated daily over a Hounsfield unit (HU) range of −1000 HU to 4000 HU. Images were reconstructed in axial mode using the manufacturer’s software (Mediso InterView FUSION, version 1.02), according to the manufacturer’s recommendations. Reconstructed images were analyzed with ImageJ (version 1.53h, NIH, USA) and CTVox (version 3.3.1, Bruker microCT) softwares to assess longitudinal changes in skeletal and metabolic parameters. All analyses were performed in a blinded manner using standardized regions of interest across time points to ensure consistency between measurements.

#### 2.2.2. Fracture count

Fractures were assessed using CT datasets acquired from NanoSPECT/CT imaging sessions. Bone structures were segmented using a global threshold of 800 HU, which provided the most accurate visualization of the skeletal architecture in this model. Three-dimensional reconstructions were examined in CTVox (version 3.3.1; Bruker microCT), allowing unrestricted 360° rotation along the X and Y axes for detailed evaluation of skeletal morphology. Fractures were identified by two independent observers in a blinded manner (MC and GM), with no knowledge of group allocation. Fractures were defined as the presence of cortical discontinuity and/or visible callus formation or deformation. All assessments were performed bilaterally for the humerus, radius, ulna, femur, tibia, and fibula. When a fracture was identified, its anatomical location was further classified as metaphyseal or diaphyseal. Metaphyseal fractures were defined as occurring within the proximal or distal one-fifth of the total bone length, whereas all other fractures were classified as diaphyseal. Inter-observer agreement was complete, with no discrepancies observed between the two evaluators.

#### 2.2.3. Abdominal adiposity

Abdominal adiposity was quantified from CT image stacks by segmenting voxels within the lumbar region (L1–L5). Adipose tissue was identified using a predefined Hounsfield unit (HU) threshold range of −50 to 200, as previously described ^(26)^. Three-dimensional regions of interest were manually delineated to include the entire abdominal compartment between L1 and L5 while excluding skeletal structures. Within this volume, voxels meeting the adipose tissue HU criteria were extracted and used to compute abdominal fat volume. All analyses were performed using consistent anatomical landmarks across animals and time points to ensure reproducibility of measurements.

#### 2.2.4. Appendicular skeleton growth

Femoral growth of the right femur was assessed by measuring femur length on three-dimensional reconstructed CT datasets using Imaris software (Version 6.4, Bitplane, Zürich, Switzerland). In cases where the right femur was fractured, measurements were performed on the contralateral femur. CT-based measurements were compared with ex vivo femur length measurements obtained using a digital caliper on specimens subsequently used for three-point bending mechanical testing. A strong correlation (R^2^ =0.91) between CT-based and caliper-based measurements confirmed the reliability of the imaging approach.

At the tissue level, longitudinal growth was evaluated in the proximal tibia. Tibiae were embedded undecalcified in polymethylmethacrylate (pMMA) and serial 7-μm sections were stained with Goldner’s trichrome. Growth plate thickness was quantified using a custom MATLAB R2025b (MathWorks, Natick, MA, USA) script. Total growth plate thickness and the thickness of the resting, proliferative, and hypertrophic zones were measured by a pathologist blinded to group allocation.

#### 2.2.5. Femur bowing

Femoral curvature was quantified from reconstructed CT images using a custom image-processing pipeline developed in MATLAB R2025b (The MathWorks, Natick, MA, USA). All measurements were scaled using the CT voxel size (0.221 mm isotropic resolution). For each animal, the central coronal slice of the right femur was selected and converted to grayscale. Images were denoised using a median filter prior to segmentation. Bone tissue was extracted from background using Otsu’s thresholding method followed by hole filling to reconstruct the medullary cavity. The femoral midline was computed by determining the medial cortical boundary and adding half of the local bone width along each row of the image. The resulting centerline was smoothed and fitted using a fourth-degree polynomial function. Femoral curvature was then derived from the fitted centerline using a geometric approach based on local tangent construction and orthogonal projections to estimate the osculating circle. The method was validated on multiple samples and showed high repeatability of curvature measurements. The radius of curvature was calculated using Euclidean distance between the estimated center of curvature and a reference point on the femoral shaft, and curvature was defined as the inverse of this radius.

### 2.3. Bone biomechanics

Whole-bone strength of right femurs was assessed by 3-point bending in accordance with published guidelines ^(27)^. Three-point bending strength was measured with a constant span length of 8 mm. Bones were tested in the antero-posterior axis with the posterior surface facing upward, centered on the support and the pressing force was applied vertically to the midshaft of the bone. Each bone was tested fully hydrated and at room temperature with a loading speed of 0.5 mm.min-1 until failure with a 500 N load cell on an Instron 5942 device (Instron, Elancourt, France) and the load-displacement curve was recorded at a 100 Hz rate by the Bluehill 3 software (Instron). Ultimate load, ultimate displacement, stiffness and work to fracture were calculated as indicated in Jepsen et al. ^(27)^. The yield load was determined as the point at which a regression line that represents a 10% loss in stiffness crosses the load-displacement curve. Post-yield displacement was computed as the displacement between yielding and fracture. Material properties were computed based on beam theory equations. Brittleness was computed as the ratio between yield strain and ultimate strain according to Bargman et al. ^(28)^.

### 2.4. Ex vivo microCT

MicroCT analyses were performed at the right femur after three-point bending with a Bruker 1272 microtomograph operated at 70 kV, 140 μA, 1000 ms integration time and imaging in ethanol 70. The isotropic pixel size was fixed at 4 μm, the rotation step at 0.25° and exposure was performed with a 0.5 mm aluminum filter. Hydroxyapatite phantoms (250 mg/cm3 and 750 mg/cm3) were used for calibration. Reconstruction of 2D projections was done with the NRecon software (Version 1.7.4.6). Prior to trabecular and cortical analyses, bones were re-axed with the Dataviewer software (version 1.6.0). A trabecular volume of interest was located 0.5 mm above the distal growth plate and extended 2 mm up. Trabecular bone was separated from cortical bone with an automatic contouring script in CTan software (Version 1.20.8.0). A cortical volume of interest (0.5 mm) was positioned 4 mm up the distal growth plate. Bone was segmented from soft tissue using global thresholding set at 300 mg/cm^3^ for trabecular bone and 700 mg/cm^3^ for cortical bone. All histomorphometrical parameters were measured with the CTan software according to guidelines and nomenclature proposed by the American Society for Bone and Mineral Research ^(29)^.

### 2.5. Fourier transform infrared imaging

After microCT analysis, the distal halves of the right femur were embedded undecalcified in polymethylmethacrylate (pMMA) following dehydration and infiltration, as previously described ^(30)^. One-micrometer-thick cross-sections of the femoral midshaft were obtained using an ultramicrotome (Leica EM UC7, Leica Microsystems, Nanterre, France) and mounted on BaF_2_ windows. Infrared spectral imaging was performed using a Bruker Hyperion II infrared microscope coupled to an Invenio-S spectrometer (Bruker Optics, Champs-sur-Marne, France) equipped with a 64 × 64 focal plane array detector. A field of view of 540 × 540 µm, corresponding to the posterior quadrant of the femoral midshaft, was analyzed. Spectra were acquired in transmission mode over the 850–2000 cm^−1^ range at 8 cm^−1^ spectral resolution, using a 15× Cassegrain objective (NA = 0.4) and 32 co-added scans per pixel. Background spectra were collected from the same BaF_2_ windows under identical conditions at the beginning and end of each experiment. A reference spectrum of pure pMMA prepared under identical conditions was used for pMMA subtraction, scaled to the intensity of the ester carbonyl band (~1730 cm^−1^). Spectral preprocessing was performed in MATLAB R2025b (The MathWorks, Natick, MA, USA) and included Mie scattering correction, pMMA subtraction, vector normalization, and Savitzky–Golay denoising (second-order polynomial, window size = 9). Second-derivative spectra were then calculated and used for curve-fitting analysis, as previously described ^(30)^. Spectral quality was verified by signal-to-noise ratio assessment in the 1850–2000 cm^−1^ region. Bone compositional parameters were calculated as follows: mineral-to-matrix ratio (integrated area of the v_1_,v_3_ phosphate bands / amide I band), mineral crystallinity/maturity (1030 cm^−1^ / 1020 cm^−1^ peak intensity ratio), carbonate-to-phosphate ratio (v_3_ carbonate band at ~1415 cm^−1^ / 1030 cm^−1^), and collagen maturity (1660 cm^−1^ / 1690 cm^−1^ intensity ratio) ^(30)^. Pixels corresponding to bone tissue were segmented using intensity-based thresholding of the v_1_,v_3_ phosphate bands, and non-bone pixels were excluded from analysis. For each parameter, pixel distributions were assessed for normality using Gaussian fitting, and all datasets were confirmed to be normally distributed (R^2^ > 0.95). Mean values for each specimen were calculated from the pixel distributions.

### 2.6. ELISA

Plasma concentrations of C-terminal telopeptide of type I collagen (CTX, ref AC-06F1, Immunodiagnostic Systems, Boldon, UK), procollagen type I N-terminal propeptide (P1NP, ref AC-33F1, Immunodiagnotic Systems), parathyroid hormone (PTH_1–84_, ref 60-2305, Quidel Corporation, Athens, OH), insulin-like growth factor 1 (IGF-1, ref MG100 Bio-Techne, Noyal Chatillonsur Seiche, France), and growth hormone (GH, ref EZRMGH-45K, Millipore, Saint-Quentin-Fallavier, France) were measured using commercially available enzyme-linked immunosorbent assays (ELISA), according to the manufacturers’ instructions. Briefly, plasma samples were thawed on ice and analyzed in duplicate. Standard curves were generated for each assay, and concentrations were calculated using a four-parameter logistic regression model. All assays were performed in a blinded manner. Samples with values below the detection limit were assigned to the lower limit of quantification for statistical analyses.

### 2.7. Statistical analysis

Sample size for mouse experiments was estimated based on previous experiments in our laboratory using fracture incidence as a primary outcome ^(25)^. Secondary outcomes of interest included effects on collagen maturity and trabecular bone mass. Statistical analyses were performed using GraphPad Prism 11.0 (GraphPad Software, La Jolla, CA, USA) unless otherwise stated. Outliers were identified using the ROUT method (Q = 1%). Homogeneity of variances was assessed using the Brown–Forsythe test. When assumptions of normality and equal variance were met, ordinary one-way ANOVA followed by Dunnett’s multiple comparisons test was performed. When normality assumptions were not met, the Kruskal– Wallis test followed by Dunn’s multiple comparisons test was used. A mixed-effects model was used to analyze changes in body weight over time, followed by Dunnett’s multiple comparisons test. Spearman correlations were performed to assess correlations between bone fracture and bone volume fraction or collagen maturity. Given that fracture number is a non-negative count variable with a positively skewed distribution (range 0–2 fractures per animal), associations between fracture number and bone parameters were assessed using Poisson regression rather than linear regression. A multivariate Poisson regression model was fitted including collagen maturity, BV/TV and CT.Ar/Tt.Ar simultaneously to assess their independent associations with fracture number, with predictors standardized (z-scores) to allow comparison of effect sizes on a common scale.

Model adequacy was assessed by evaluating the dispersion parameter; a value close to 1 confirmed that the variance-to-mean assumption of the Poisson distribution was appropriate and that a negative binomial model was not required. Results are reported as incidence rate ratios (IRR) with 95% confidence intervals, derived from the exponentiated regression coefficients. Statistical analyses were performed in MATLAB (version R2025b) using the fitglm function, and significance was set at p < 0.05. Unless otherwise stated, data are presented as mean ± SD. Differences were considered statistically significant at p<0.05.

## 3. Results

### 3.1. Effects of GL-0001 on body weight and abdominal adiposity

First, we evaluated whether zoledronic acid or GL-0001 influenced body weight, as reductions in body weight have been consistently reported with dual GIP/GLP-1 receptor agonists, another class of incretin-based therapies ^(31)^. As expected, at 5 weeks of age, OI+saline animals exhibited a significantly lower body weight compared with WT+saline animals (−27%, p<0.001) (Figure 1A). However, no significant differences in body weight were observed among the three OI groups at treatment initiation. Over the course of the study, body weight increased in all groups. Although zoledronic acid- and GL-0001-treated OI animals tended to display greater weight gain compared with OI+saline animals, neither treatment significantly altered body weight gain during the study period (Figures 1B-C).

**Figure 1.**
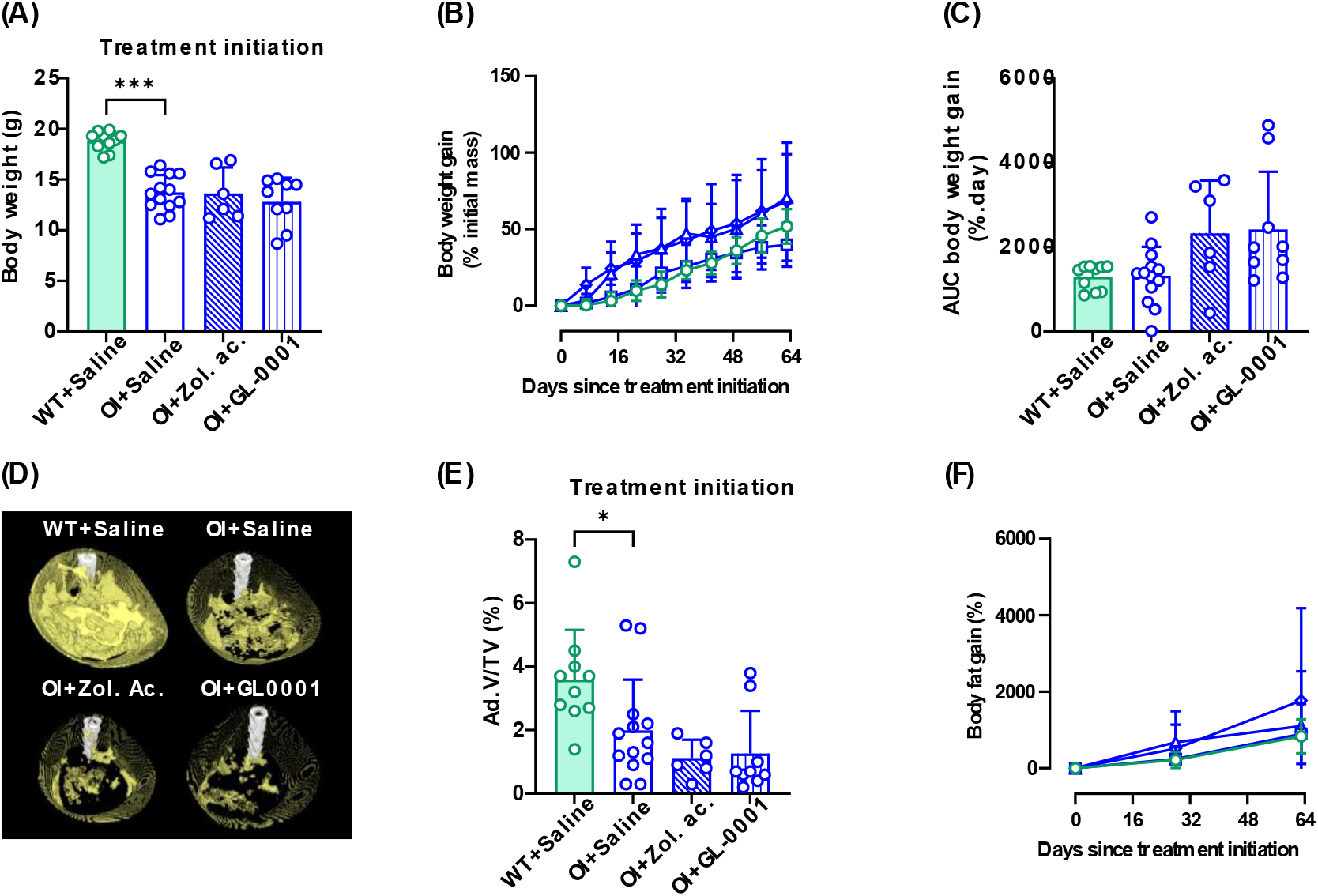
Effects of zoledronic acid and GL-0001 on body weight and abdominal adiposity. Animals were weighed weekly, and abdominal adiposity was assessed at the end of the 9-week treatment period. (A) Body weight at treatment initiation; (B) body weight gain over the study period; (C) area under the curve (AUC) of body weight gain; (D) representative three-dimensional reconstructions of abdominal adipose tissue derived from SPECT scans, with adipose tissue shown in yellow and the spine in white; (E) abdominal adipose volume fraction at treatment initiation; and (F) change in abdominal adiposity over the study period. WT + saline animals are represented by solid green bars or open circles, OI + saline animals by open bars with blue outlines or open squares, OI + zoledronic acid animals by blue diagonal-hatched bars or open triangles, and OI + GL-0001 animals by blue vertical-hatched bars or open diamonds. Data are presented as mean ± SD. Statistical analyses were performed using one-way ANOVA followed by Dunnett’s multiple comparisons test. *p<0.05, **p<0.01, and ***p<0.001 versus OI+saline. n=10 for WT+saline, n=13 for OI+saline, n=6 for OI+zoledronic acid, and n=9 for OI+GL-0001.

Given the previously reported role of the GIP/GIPr pathway in adipose tissue regulation ^(32)^, abdominal fat percentage was assessed as a surrogate marker of adiposity ^(33)^. At treatment initiation, OI+saline animals displayed significantly lower abdominal adiposity compared with WT+saline animals (−45%, p=0.032) (Figures 1D-E). No significant differences were observed among the three OI groups at baseline. Furthermore, neither zoledronic acid nor GL-0001 treatment significantly modified abdominal adiposity throughout the study (Figure 1F).

### 3.2. Effects of GL-0001 on appendicular and axial growth

Because type III OI is characterized by impaired longitudinal growth and short stature, we evaluated the effects of zoledronic acid and GL-0001 on femoral growth. Femur length measured by digital caliper was strongly correlated with measurements obtained from micro-CT analyses (R^2^=0.91, p<0.001). At baseline, OI animals exhibited significantly shorter femora than WT animals (12.2 ± 0.6 mm vs. 13.5 ± 0.2 mm, p<0.001) (Figure 2A). No differences in femur length were observed among OI groups before treatment initiation, indicating comparable skeletal growth at study entry. Following treatment, GL-0001 increased femur growth by 5% (p=0.016) (Figure 2B). On the other hand, the growth speed, expressed as micrometers change in femur length per day, was augmented by 36% in GL-0001-treated animals as compared with saline-treated OI animals (p=0.002) (Figure 2C). Zoledronic acid had no significant effect. Because long-bone bowing is a common feature of type III OI, we next assessed whether the increase in femur length observed with GL-0001 was associated with changes in bone curvature. Contrary to expectations, femoral curvature did not differ between saline-treated OI and WT animals (Figure 2D). Consistent with this finding, GL-0001 did not alter femoral curvature. In contrast, zoledronic acid significantly increased femoral bowing by 89% compared with saline-treated OI animals (p<0.001). To investigate potential mechanisms underlying the effects of GL-0001 on longitudinal growth, growth plate morphology was assessed in the proximal tibia. GL-0001 treatment increased total growth plate thickness by 22% compared with saline-treated OI animals (p=0.038; Figure 2E). Analysis of individual growth plate compartments revealed that the proliferative zone was 23% thinner in OI+saline animals than in WT animals (p=0.031) (Supplementary Figure 2). In contrast, GL-0001 treatment resulted in a selective expansion of the proliferative zone, which was 52% thicker than in saline-treated controls (p<0.001). Zoledronic acid had no effect on total growth plate thickness or on the thickness of individual growth plate zones. Circulating levels of growth hormone (GH) and insulin-like growth factor-1 (IGF-1) were subsequently quantified (Figures 2F-G). Serum GH concentrations were significantly lower in WT animals than in saline-treated OI animals (−90%, p=0.018). Neither zoledronic acid nor GL-0001 affected circulating GH levels. In contrast, serum IGF-1 levels were 27% higher in WT animals than in saline-treated OI animals (p<0.001). However, neither treatment significantly affected circulating IGF-1 concentrations.

**Figure 2.**
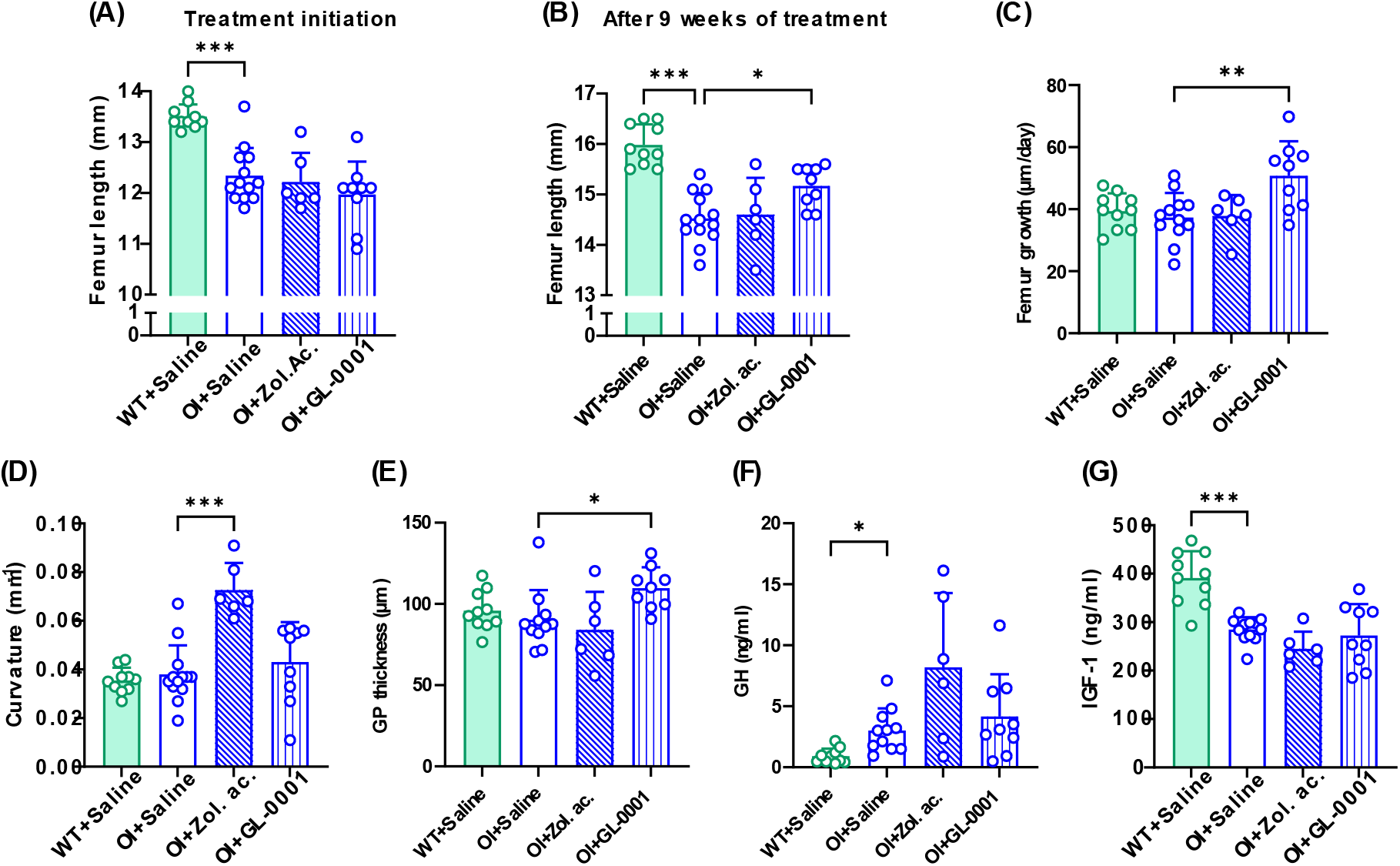
Effects of zoledronic acid and GL-0001 on longitudinal bone growth and growth plate morphology. Femur length was measured at treatment initiation (A) and after 9 weeks of treatment (B), and longitudinal growth velocity was subsequently determined (C). Given the prevalence of long-bone bowing in type III OI, femoral curvature was also assessed (D). Growth plate thickness was evaluated histologically (E) and circulating concentrations of growth hormone (GH) (F) and insulin-like growth factor-1 (IGF-1) (G) were measured. WT+saline animals are represented by solid green bars, OI+saline animals by open bars with blue outlines, OI+zoledronic acid animals by blue diagonal-hatched bars and OI+GL-0001 animals by blue vertical-hatched bars. Data are presented as mean ± SD. Statistical analyses were performed using one-way ANOVA followed by Dunnett’s multiple comparisons test. *p<0.05, **p<0.01, and ***p<0.001 versus OI+saline. n=10 for WT+saline, n=11-13 for OI+saline, n=6 for OI+zoledronic acid, and n=9 for OI+GL-0001. Differences between the planned and analyzed sample sizes in OI+saline animals for circulating GH and IGF-1 levels resulted from exclusion of outlier values identified by the ROUT method.

### 3.3. Effects of GL-0001 on fracture incidence

We next evaluated the effects of saline, zoledronic acid, or GL-0001 treatment on fracture incidence throughout the study period. Due to the voxel resolution of the SPECT/CT acquisitions, only fractures affecting the appendicular skeleton were quantified. WT animals did not exhibit fractures either at treatment initiation or during follow-up (Figure 3A). In contrast, OI animals presented with a significantly higher spontaneous fracture burden at baseline, with an average of 2 ± 1 fractures per animal (p<0.001). Fractures were predominantly characterized by mineralized calluses, followed by non-unions and, to a lesser extent, bone deformities (Figure 3B). Fractures were mainly localized to diaphysis (66%), whereas metaphyseal fractures accounted for 34% of cases. No significant differences in fracture number, fracture type, or fracture localization were observed among the three OI groups at treatment initiation, indicating comparable skeletal fragility prior to treatment onset. Interestingly, GL-0001 treatment reduced the incidence of newly occurring fractures by 90% during the 9-week treatment period (p=0.004) (Figure 3C). In contrast, although zoledronic acid treatment showed a trend toward reduced fracture incidence, this effect did not reach statistical significance (p=0.509).

**Figure 3.**
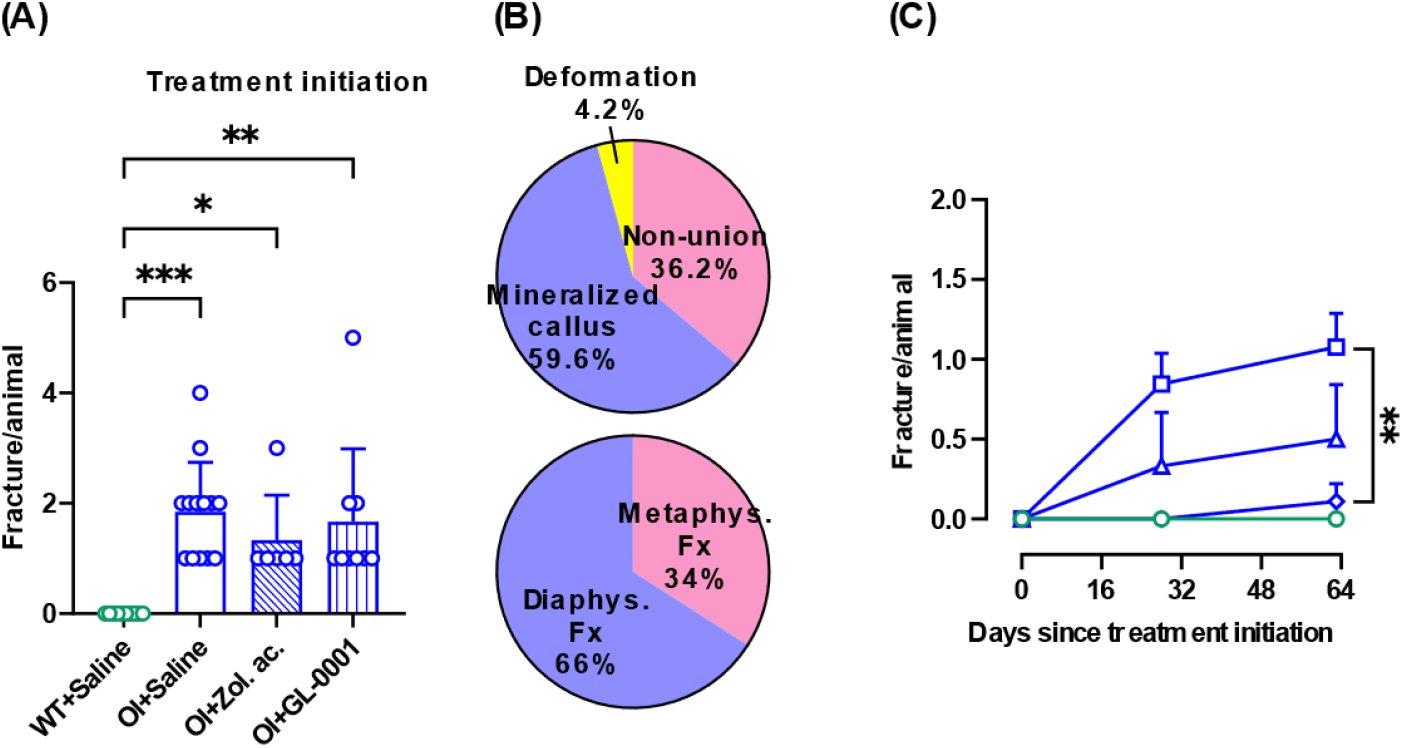
Effects of zoledronic acid and GL-0001 on fracture occurrence. The number of long bone fractures per animal was determined at treatment initiation (A). The type and anatomical localization of fractures (B) were also recorded. The number of newly occurring fractures during the treatment period was quantified (C). WT + saline animals are represented by solid green bars or open circles, OI + saline animals by open bars with blue outlines or open squares, OI + zoledronic acid animals by blue diagonal-hatched bars or open triangles, and OI + GL-0001 animals by blue vertical-hatched bars or open diamonds. Data are presented as mean ± SD. Statistical analyses were performed using one-way ANOVA followed by Dunnett’s multiple comparisons test. *P < 0.05, **P < 0.01, and ***P < 0.001 versus OI + saline. n = 10 for WT + saline, n = 13 for OI + saline, n = 6 for OI + zoledronic acid, and n = 9 for OI + GL-0001.

### 3.4. Effects of GL-0001 on bone biomechanics

The effects of 9 weeks of saline, zoledronic acid, or GL-0001 treatment on bone mechanical properties were assessed by three-point bending (Table 1). As expected, OI+saline mice exhibited marked impairments in structural properties compared with WT+saline animals, including a 59% decrease in ultimate load (p<0.001), a 60% decrease in yield load (p<0.001), a 65% decrease in stiffness (p<0.001), a 75% decrease in post-yield displacement (p<0.001), and an 82% decrease in work-to-fracture (p<0.001). At the material level, OI+saline mice also showed a 27% reduction in elastic modulus (p=0.006) and a 76% reduction in post-yield strain (p<0.001), together with a 125% increase in brittleness (p<0.001). At the structural level, zoledronic acid showed trends toward increased ultimate load (26%, p=0.093), yield load (31%, p=0.105), and stiffness (45%, p=0.057), although these differences did not reach statistical significance. No significant effects of zoledronic acid were observed for material-level parameters. GL-0001 treatment also showed a trend toward increased ultimate load (23%, p=0.119). At the material level, GL-0001 significantly increased ultimate stress (26%, p=0.022) and yield stress (30%, p=0.014), restoring values toward those observed in WT animals. In addition, bone ductility was improved, as reflected by a 32% reduction in brittleness (p=0.002).

**Table 1:** Structural and material properties of femora determined by three-point bending testing.

|  | WT+Saline | OI+Saline | OI+Zol. Ac. | OI+GL-0001 |
| --- | --- | --- | --- | --- |
| **Structural parameters** |  |  |  |  |
| Ultimate load (N) | 17.8 ± 2.3 *** | 7.3 ± 1.1 | 9.2 ± 1.9 | 9.0 ± 0.3 |
| Yield load (N) | 13.8 ± 2.0 *** | 5.5 ± 0.9 | 7.2 ± 1.6 | 6.9 ± 1.1 |
| Stiffness (N.mm<sup>-1</sup>) | 95 ± 15 *** | 33 ± 7 | 48 ± 13 | 42 ± 12 |
| Post-yield displacement (mm) | 0.44 ± 0.20 *** | 0.11 ± 0.05 | 0.09 ± 0.04 | 0.18 ± 0.06 |
| Work-to-fracture (N.mm) | 7.6 ± 2.3 *** | 1.4 ± 0.6 | 1.3 ± 0.4 | 1.7 ± 0.3 |
| **Material parameters** |  |  |  |  |
| Ultimate stress (MPa) | 88 ± 9 | 76 ± 15 | 68 ± 14 | 93 ± 15 * |
| Yield stress (MPa) | 69 ± 10 | 57 ± 13 | 52 ± 9 | 74 ± 12 * |
| Elastic modulus (GPa) | 6.3 ± 0.8 ** | 4.6 ± 0.9 | 4.5 ± 1.1 | 6.6 ± 2.2 |
| Post-yield strain | 0.033 ± 0.016<br>*** | 0.008 ± 0.003 | 0.006 ± 0.003 | 0.013 ± 0.004 |
| Brittleness (%) | 28 ± 11 *** | 63 ± 12 | 64 ± 10 | 43 ± 8 ** |

Data are presented as mean ± SD. Statistical analyses were performed using one-way ANOVA followed by Dunnett's multiple-comparisons test, except for post-yield displacement, work-to-fracture, elastic modulus, and post-yield strain, which were analyzed using the Kruskal–Wallis test followed by Dunn's multiple-comparisons test. \*p < 0.05, \*\*p < 0.01, \*\*\*p < 0.001 versus OI+Saline. n = 10 animals/group for the WT+Saline and OI+Saline groups, n = 6 for the OI+Zoledronic Acid group, and n = 8 for the OI+GL-0001 group. Differences between the planned and analyzed sample sizes were due to the presence of fracture calluses at the femoral diaphysis, which precluded mechanical testing.

### 3.5. Effects of GL-0001 on bone microstructure and turnover

Trabecular and cortical bone microarchitecture were assessed by ex vivo micro-computed tomography. Analysis of the distal femur revealed marked alterations in trabecular bone microarchitecture in OI+saline animals compared with WT+saline controls (Figure 4A), including reductions in bone volume fraction (BV/TV; −57%, p<0.001), trabecular number (Tb.N; −49%, p<0.001), and trabecular thickness (Tb.Th; −13%, p<0.001), together with an increase in trabecular separation (Tb.Sp; 26%, p=0.003) (Figures 4B-E). Zoledronic acid treatment significantly improved trabecular bone microarchitecture, as evidenced by increases in BV/TV (213%, p<0.001) and Tb.N (230%, p<0.001), as well as a reduction in Tb.Sp (−27%, p<0.001). In contrast, GL-0001 treatment for 9 weeks did not significantly affect trabecular bone microarchitecture.

**Figure 4.**
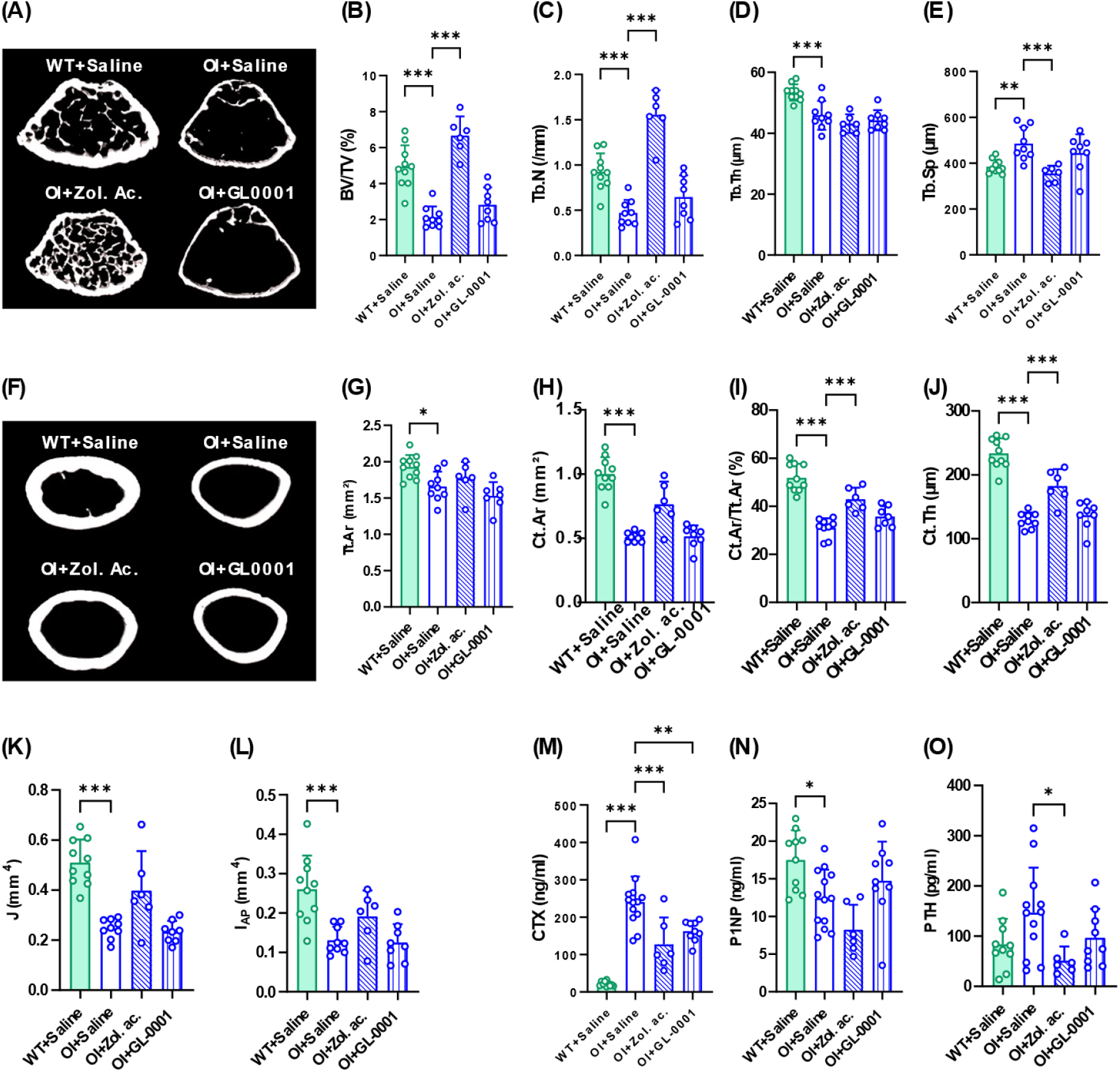
Effects of zoledronic acid and GL-0001 on trabecular and cortical microarchitecture and circulating bone remodeling markers. Trabecular and cortical bone microarchitecture were assessed in the distal femoral metaphysis and diaphysis using ex vivo micro-computed tomography (μCT). (A) Three-dimensional reconstructions of the distal femoral metaphysis; (B–E) trabecular microarchitecture parameters; (F) three-dimensional reconstructions of the femoral diaphysis; and (G–L) cortical bone microarchitecture parameters. Circulating levels of the bone remodeling markers CTX (M), P1NP (N), and PTH (O) were also measured. WT + saline animals are represented by solid green bars, OI + saline animals by open bars with blue outlines, OI + zoledronic acid animals by blue diagonal-hatched bars, and OI + GL-0001 animals by blue vertical-hatched bars. Data are presented as mean ± SD. Statistical analyses were performed using one-way ANOVA followed by Dunnett’s multiple comparisons test. *P < 0.05, **P < 0.01, and ***P < 0.001 versus OI + saline. n=10 for WT+saline, n=8-9 for OI+saline, n=6 for OI+zoledronic acid, and n=78 for OI+GL-0001. Differences between the planned and analyzed sample sizes were due to the presence of fracture calluses at the femoral diaphysis, which precluded microstructural investigations or resulted from exclusion of outlier values identified by the ROUT method.

Because three-point bending predominantly reflects cortical bone mechanical properties, cortical bone geometry was subsequently evaluated (Figure 4F). Compared with WT+saline animals, OI+saline mice exhibited lower total cross-sectional area (Tt.Ar; −14%, p=0.015), cortical area (Ct.Ar; −49%, p<0.001), cortical area fraction (Ct.Ar/Tt.Ar; −40%, p<0.001), cortical thickness (Ct.Th; −45%, p<0.001), polar moment of inertia (J; −52%, p<0.001), and anteroposterior moment of inertia (I_AP_; −50%, p<0.001) (Figures 4G-L). Zoledronic acid significantly increased Ct.Ar/Tt.Ar (38%, p<0.001) and Ct.Th (42%, p<0.001), whereas no significant changes were observed for the remaining cortical parameters. GL-0001 treatment did not significantly alter cortical bone geometry.

Circulating markers of bone remodeling were subsequently quantified (Figures 4M-O). Compared with WT+saline animals, OI+saline mice exhibited markedly elevated serum CTX levels (+1,003%, p<0.001) and reduced serum P1NP levels (−29%, p=0.018), whereas circulating PTH concentrations did not differ between groups. Zoledronic acid treatment significantly reduced serum CTX levels by 47% compared with saline-treated OI animals (p<0.001) but did not alter circulating P1NP concentrations. In contrast, serum PTH levels were reduced by 65% following zoledronic acid treatment (p=0.018). Similarly, daily administration of GL-0001 for 9 weeks significantly decreased serum CTX levels by 32% (p=0.004) without affecting circulating P1NP or PTH concentrations.

### 3.6. Effects of GL-0001 on bone material properties

Bone material properties were subsequently evaluated to determine whether the effects of GL-0001 on fracture incidence and bone mechanical properties were associated with alterations in tissue composition (Figure 5 and supplementary Table 1). Compared with WT+saline animals, OI+saline mice exhibited lower collagen maturity (−13%, p < 0.001) and higher mineral-to-matrix ratio (+37%, p<0.001), acid phosphate content (+7%, p=0.003), and nanoporosity (+45%, p<0.001). Zoledronic acid treatment did not significantly affect any of the assessed bone material properties. In contrast, GL-0001 significantly increased collagen maturity by 12% (p<0.001) and showed a trend toward a reduction in the mineral-to-matrix ratio (p=0.092).

**Figure 5.**
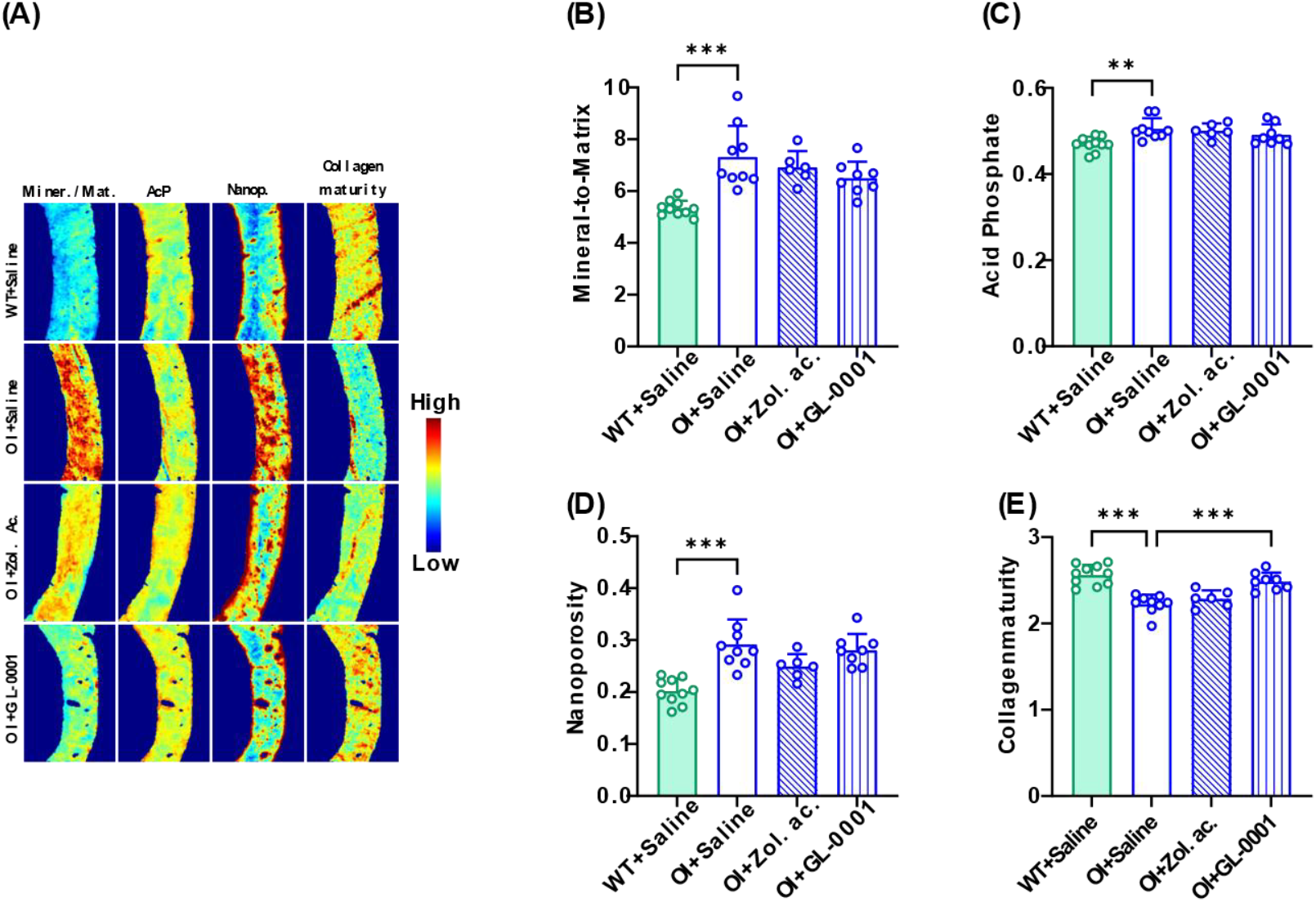
Effects of zoledronic acid and GL-0001 on bone material properties. Bone material properties were assessed by Fourier transform infrared imaging (FTIRI) after 9 weeks of treatment. (A) Representative FTIRI maps of bone material parameters; quantitative analyses of (B) mineral-to-matrix ratio, (C) acid phosphate content, (D) nanoporosity, and (E) collagen maturity. WT + saline animals are represented by solid green bars, OI + saline animals by open bars with blue outlines, OI + zoledronic acid animals by blue diagonal-hatched bars, and OI + GL-0001 animals by blue vertical-hatched bars. Data are presented as mean ± SD. Statistical analyses were performed using one-way ANOVA followed by Dunnett’s multiple comparisons test. *P < 0.05, **P < 0.01, and ***P < 0.001 versus OI + saline. n=10 for WT+saline, n=9 for OI+saline, n=6 for OI+zoledronic acid, and n=8 for OI+GL-0001. Differences between the planned and analyzed sample sizes were due to the presence of fracture calluses at the femoral diaphysis, which precluded investigations of bone material properties.

### 3.7. Regression analysis between fracture, bone volume fraction and collagen maturity

Spearman correlation analysis showed no association between the number of fractures and either BV/TV (r=0.037, p=0.840) or Ct.Ar/Tt.Ar (r=-0.214, p=0.239).In contrast, collagen maturity was significantly associated with fracture number (r=-0.394, p=0.023), indicating that animals with higher collagen maturity exhibited fewer fractures. To further assess these associations while accounting for potential confounding between predictors, multivariate Poisson regression analysis was performed including collagen maturity, BV/TV, and Ct.Ar/Tt.Ar as independent variables (Table 2). Model fit was appropriate for a Poisson distribution, with an estimated dispersion parameter close to 1 (1.02), indicating no overdispersion. Only collagen maturity remained independently associated with fracture burden (IRR=0.49, 95% CI [0.25–0.94], p=0.032), whereas neither BV/TV (IRR=1.05, p=0.857) nor Ct.Ar/Tt.Ar (IRR=1.25, p=0.386) showed a significant association with fracture number after adjustment. These findings suggest that matrix quality, rather than bone mass or cortical area fraction, is the main determinant of fracture susceptibility in this model.

**Table 2:** Multivariate Poisson regression analysis.

| Variable | $\beta$ | SE | IRR [95% CI] | p value |
| --- | --- | --- | --- | --- |
| Collagen maturity | -0.716 | 0.334 | 0.49 [0.25-0.94] | 0.032 |
| BV/TV | 0.049 | 0.274 | 1.05 [0.61-1.80] | 0.857 |
| Ct.Ar/Tt.Ar | 0.222 | 0.257 | 1.25 [0.76-2.07] | 0.386 |

SE: Standard error, IRR: Incidence rate ratio, 95% CI: 95% confidence interval

## 4. Discussion

In this study, we demonstrate that GL-0001, a novel unimolecular dual GIP/GLP-2 analog, markedly improves skeletal fragility in the oim/oim mouse model of severe osteogenesis imperfecta. The most striking finding is a profound reduction in spontaneous fracture incidence, accompanied by improvements in bone mechanical competence and material-level properties, despite minimal effects on bone microarchitecture. In contrast, zoledronic acid primarily improved bone structure but did not significantly affect fracture incidence or bone material properties. Together, these findings suggest that GL-0001 enhances bone strength predominantly through qualitative improvements in bone tissue rather than quantitative changes in bone mass.

An important finding of the present study is the dissociation between improvements in bone quantity and fracture protection. Although bisphosphonate treatment significantly increased BV/TV and Ct.Ar/Tt.Ar, these structural improvements were not accompanied by a significant reduction in fracture number. In contrast, peptide treatment significantly reduced fracture burden while primarily increasing collagen maturity. Consistent with these group-specific effects, Poisson regression analysis identified collagen maturity as the only bone parameter significantly associated with fracture number, whereas BV/TV and Ct.Ar/Tt.Ar were not associated with fracture burden. Collectively, these findings support the hypothesis that improvements in collagen matrix quality contribute to the antifracture efficacy of the peptide and emphasize that bone quality, beyond bone quantity alone, is an important determinant of skeletal fragility in this model.

A notable finding of this study is the increase in femur length in GL-0001–treated oim/oim mice, associated with a selective expansion of the proliferative zone of the growth plate. Impaired longitudinal growth is a hallmark of severe OI, particularly in the oim/oim model, which recapitulates growth retardation and skeletal fragility observed in patients with severe form of the disease. The expansion of the proliferative zone suggests that GL-0001 enhances chondrocyte proliferation and/or survival. Importantly, these effects occurred without changes in circulating growth hormone (GH) or insulin-like growth factor-1 (IGF-1), indicating that GL-0001 does not act through systemic activation of the classical somatotropic axis. Analysis of publicly available dataset of gene expression in the Aga2 OI mice reported the presence of *Gipr* and *Glp2r* expression in the growth plate chondrocyte ^(34)^. This supports the hypothesis of a local action at the growth plate.

The most clinically relevant outcome of this study is the 90% reduction in fracture incidence in GL-0001-treated animals. This effect occurred in the absence of major improvements in trabecular or cortical microarchitecture, highlighting a dissociation between bone structure and bone strength. This observation underscores the importance of bone quality, particularly in OI, where fragility is primarily driven by qualitative defects in type I collagen rather than bone mass per se ^(4)^. In the oim/oim model, abnormal collagen assembly leads to altered mineralization patterns and reduced tissue toughness, which cannot be fully captured by microarchitectural indices alone ^(35)^. Previously, we also evidenced a defaut in enzymatic collagen cross-linking in young oim females ^(7)^. Thus, the reduction in fracture burden observed with GL-0001 likely reflects improvements in tissue-level properties governing crack initiation and propagation rather than structural reinforcement. At the tissue level, GL-0001 significantly increased ultimate and yield stress, improved ductility, and reduced brittleness. These changes were associated with increased collagen maturity, a surrogate marker of enzymatic collagen crosslinking ^(30)^, and a trend toward normalization of the mineral-to-matrix ratio. This mechanism of action is aligned with the strategic design of dual GIP/GLP-2 analogues that have been designed to increase Lox expression and improve enzymatic collagen cross-linking ^(23)^. In a previous animal model of ovariectomy-induced bone fragility, GL-0001 was already proven efficient in enhancing bone strength by ameliorating collagen maturity and normalizing mineral-to-matrix ratio ^(23)^. Given that OI is fundamentally a collagen disorder caused by mutations affecting type I collagen structure and processing, even modest improvements in collagen maturation can have disproportionate biomechanical consequences.

As expected, zoledronic acid improved trabecular microarchitecture and reduced serum CTX levels, consistent with its potent antiresorptive action ^(36)^. However, these structural improvements were not accompanied by changes in bone material properties or fracture incidence in this study. This dissociation highlights a key limitation of repositioning osteoporotic drugs in OI: while increasing bone mass may partially improve structural parameters, it does not correct the underlying collagen matrix defects that determine tissue fragility. In rodents, but also in humans, the efficacy of bisphosphonates in reducing fracture occurrence in OI remains controversial ^(8,37,38)^. Recent clinical studies in OI patients have similarly shown that Setrusumab, an anti-sclerostin monoclonal antibody, or the anabolic sequence of Teriparatide followed by Zoledronic acid, although increase bone mineral density but without fully normalizing fracture risk, suggesting that improving bone mineral density and mass might not be the target in OI ^(12,13)^.

Both GL-0001 and zoledronic acid reduced serum CTX levels, indicating a decrease in bone resorption. In contrast, circulating P1NP levels were unchanged across treatment groups, suggesting that neither intervention had a major impact on bone formation over the study period. Zoledronic acid, but not GL-0001, significantly reduced circulating PTH concentrations, consistent with its well-established effects on calcium homeostasis and bone turnover suppression. Overall, the pattern observed with GL-0001 is in line with previous findings in ovariectomized models, where similar modulation of bone resorption markers was reported without marked effects on bone formation or endocrine regulators ^(23)^. Importantly, GL-0001 did not alter circulating GH, IGF-1, or PTH levels, further supporting the concept that its skeletal effects are unlikely to be mediated through systemic endocrine pathways, but rather through local actions within bone tissue.

This study has several limitations. First, the relatively short treatment duration may have limited our ability to detect longer-term effects on cortical remodeling and bone geometry, which typically require more extended periods to fully manifest. Second, the multivariate Poisson regression analysis was conducted in a relatively small cohort with a low number of fracture events, which may limit the stability of the estimates and increase the risk of overfitting. Consequently, these findings should be considered hypothesis-generating and warrant validation in larger independent studies. Third, the present study was conducted exclusively in female mice; given potential sex-specific differences in bone phenotype ^(6,24)^, validation of these findings in male animals will be important to ensure broader translational relevance. Future studies should aim to dissect the receptor-specific mechanisms underlying GL-0001 action, with particular emphasis on osteoblast and chondrocyte biology, and to delineate downstream signaling pathways involved in collagen maturation and extracellular matrix organization. In addition, longer-term studies will be required to determine whether the improvements in bone material properties and reduced fracture incidence observed here are sustained over time and translate into durable protection against skeletal fragility.

In conclusion, GL-0001 significantly improves skeletal fragility in a severe model of osteogenesis imperfecta through mechanisms primarily involving enhancement of bone material properties and collagen quality, rather than increases in bone mass or microarchitecture. These findings highlight the therapeutic potential of targeting bone quality in disorders such as OI, where collagen abnormalities are central to disease pathophysiology.

## 5. Author contributions

Conceptualization: G. Mabilleau & C. Behets, Formal analysis: M. Couchot, A. Mieczkowska, Investigation: M. Couchot, A. Chrétien & A. Mieczkowska, Resources: G. Mabilleau, Data curation: G. Mabilleau, Writing – original draft: G. Mabilleau, Writing – Review and editing: M. Couchot, A. Chrétien, A. Mieczkowska & C. Behets, Funding acquisition: G. Mabilleau

## 6. Acknowledgements

The authors thank the HiMolA platform – Inserm UMR_S 1229/SFR ICAT 4208 – University of Angers for support with bone assessment. The SCIAM - SFR ICAT 4208 – University of Angers platform is also thanked for the use of the Imaris software. This work was supported by an institutional grant from SATT Ouest Valorisation (#DV2541) to GM.

## 7. Data availability statement

The data that support the findings of this study are available from the corresponding author upon reasonable request.

## 9. Ethics approval statement

All procedures were conducted in accordance with the European Union Directive 2010/63/EU for animal experiments and the Belgian federal regulations for animal experimentation and were approved by the Institutional Animal Care and Use Committee of the Université catholique de Louvain (protocol #2020/UCL/MD/02).

