## Supplementary data for "GL-0001, a first-in-class dual GIP/GLP-2 analogue, reduces bone fragility by enhancing bone material properties in a murine model of type III osteogenesis imperfecta"

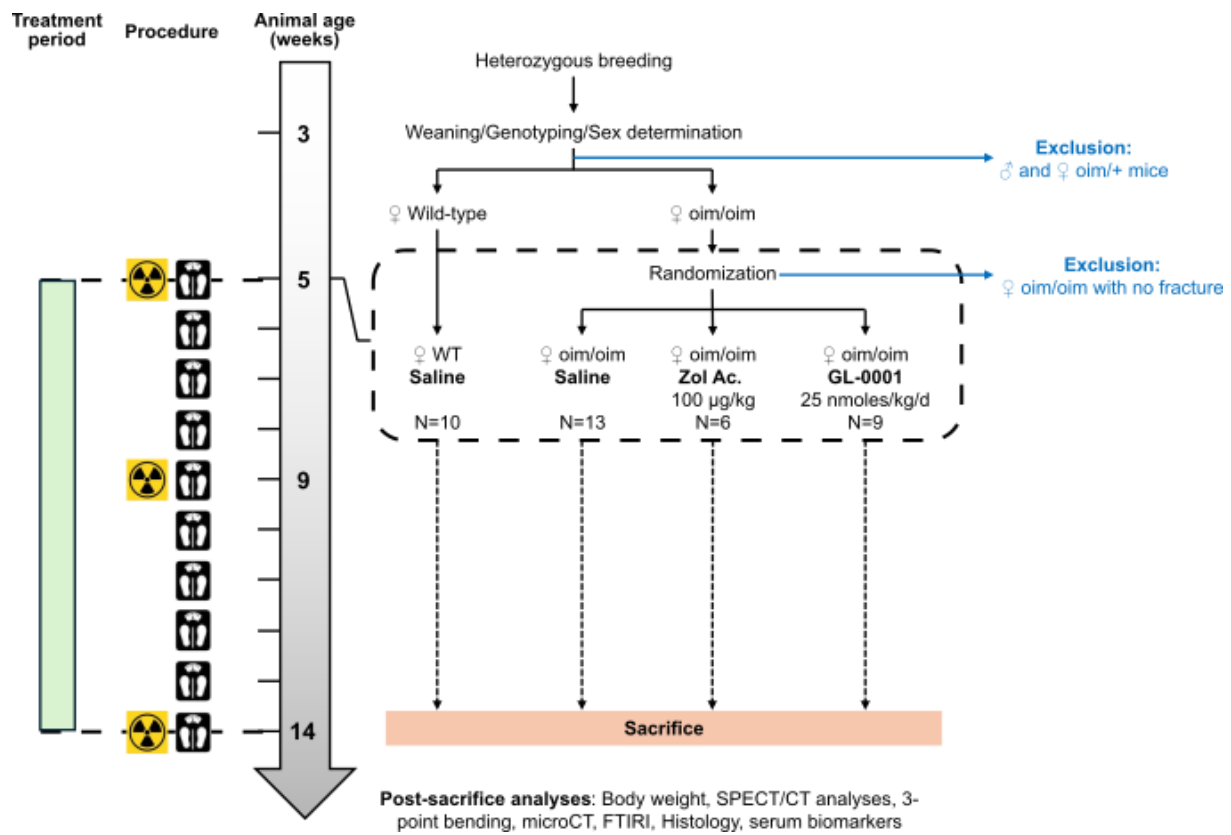

**Supplementary Figure 1 : Experimental design**

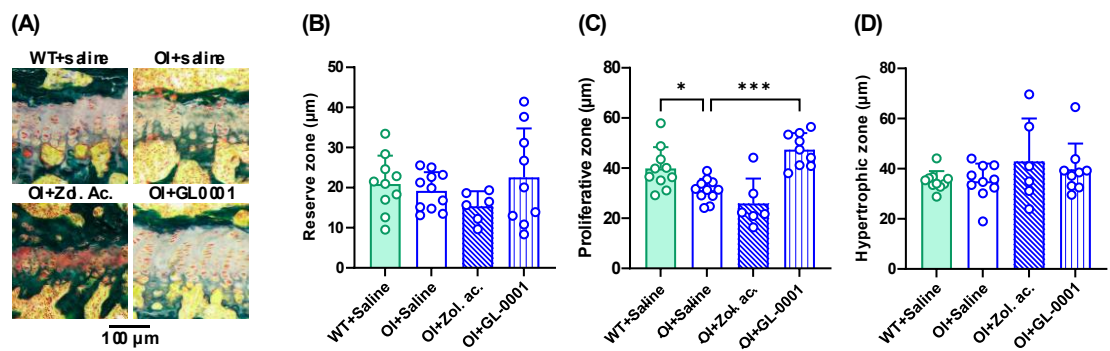

**Supplementary Figure 2.** Growth plate histology and zonal thickness. (A) Representative histological images of the growth plate stained with Goldner's trichrome. Quantification of the thickness of the (B) reserve zone, (C) proliferative zone, and (D) hypertrophic zone of the growth plate. WT+saline animals are represented by solid green bars, OI+saline animals by open bars with blue outlines, OI+zoledronic acid animals by blue diagonal-hatched bars and OI+GL-0001 animals by blue vertical-hatched bars. Data are presented as mean ± SD. Statistical analyses were performed using one-way ANOVA followed by Dunnett's multiple comparisons test. \*$p < 0.05$ and \*\*\*$p < 0.001$ versus OI+saline. $n = 10$ for WT+saline, $n = 11-13$ for OI+saline, $n = 6$ for OI+zoledronic acid, and $n = 9$ for OI+GL-0001.

**Supplementary Table 1: Bone material properties of feur diaphysis determined by Fourier transform infrared imaging (FTIRI)**

|  | WT+Saline | OI+Saline | OI+Zol. Ac. | OI+GL-0001 |
| --- | --- | --- | --- | --- |
| Mineral-to-Matrix | 5.3 ± 0.3 *** | 7.3 ± 1.2 | 6.9 ± 0.6 | 6.5 ± 0.6 |
| Mineral crystallinity | 1.28 ± 0.05 | 1.27 ± 0.02 | 1.27 ± 0.02 | 1.28 ± 0.02 |
| Crystal size index | 0.871 ± 0.017 | 0.869 ± 0.015 | 0.866 ± 0.008 | 0.867 ± 0.014 |
| Carb./Phos. | 0.259 ± 0.008 | 0.267 ± 0.027 | 0.260 ± 0.016 | 0.250 ± 0.014 |
| Acid Phosphate | 0.470 ± 0.017 ** | 0.505 ± 0.025 | 0.501 ± 0.017 | 0.492 ± 0.024 |
| Nanoporosity | 0.202 ± 0.024 *** | 0.292 ± 0.048 | 0.249 ± 0.024 | 0.281 ± 0.031 |
| Collagen Maturity | 2.56 ± 0.12 *** | 2.22 ± 0.11 | 2.29 ± 0.10 | 2.49 ± 0.10 *** |

Statistical analyses were performed with one way ANOVA with Dunnett's multiple comparisons test. \*: p<0.05; \*\*: p<0.01; \*\*\*: p<0.001 vs. OI+Saline. n = 10 animals/group for the WT+Saline group, n = 9 for the OI+Saline group, n = 6 for the OI+Zoledronic Acid group, and n = 8 for the OI+GL-0001 group. Differences between the planned and analyzed sample sizes were due to the presence of fracture calluses at the femoral diaphysis, which precluded FTIRI investigations.
